# Multicellular PI control for gene regulation in microbial consortia

**DOI:** 10.1101/2022.03.21.485171

**Authors:** Vittoria Martinelli, Davide Salzano, Davide Fiore, Mario di Bernardo

**Affiliations:** Department of Electrical Engineering and Information Technology, University of Naples Federico II, Via Claudio 21, 80125 Naples, Italy; SSM - School for Advanced Studies, Naples, Italy; Department of Mathematics and Applications “R. Caccioppoli”, University of Naples Federico II, Via Cintia, Monte S.Angelo, 80126 Naples, Italy

## Abstract

We describe two multicellular implementations of the classical P and PI feedback controllers for the regulation of gene expression in a target cell population. Specifically, we propose to distribute the proportional and integral actions over two different cellular populations in a microbial consortium comprising a third target population whose output needs to be regulated. By engineering communication among the different cellular populations via appropriate orthogonal quorum sensing molecules, we are able to close the feedback loop across the consortium. We derive analytical conditions on the biological parameters guaranteeing the regulation of the output of the target population and we validate the robustness and modularity of proposed control schemes via *in silico* experiments in BSim, a realistic agent-based simulator of bacterial populations.

## I. Introduction

Synthetic Biology aims at the design and implementation of novel, and more reliable, genetic circuits by employing engineering principles, with applications spanning different fields. Examples include the production of sustainable biofuels, the design of biosensors able to detect the presence of environmental pollutants, e.g., [1], or in medicine, for the treatment and prevention of infections and other diseases [2]. However, interactions of synthetic circuits with the host cell or with other genetic circuits, as well as unavoidable nonlinear and stochastic effects, may cause problems such as poor modularity and undesired behavior [3]. A possible solution to tackle these problems is to embed in the cells engineered feedback mechanisms to achieve more stable and robust operation of the genetic circuits of interest in a variety of operating conditions. In particular, this could facilitate the transition from highly controlled laboratory conditions to practical real-world applications [4].

Indeed, several solutions have been proposed in the literature in this direction, such as the antithetic feedback controller guaranteeing robust perfect adaptation in noisy biomolecular networks [5], [6], or the implementation via biological molecules in a single cell of a proportional-integral-derivative control strategy [7]. Further examples of synthetic feedback mechanisms embedded in single cells can be found in [8], [9].

The use of biomolecular PID controllers is particularly appealing as it allows achieving perfect robust adaptation via the integral action as well as to exploit the proportional and derivative actions to modulate the steady-state and transient dynamics of the controlled process. However, embedding all the required circuits to implement a PID controller in a single cell could cause excessive metabolic burden and be cumbersome to implement *in vivo*; also requiring a complete re-design if the target process to be regulated changes or the parameters of the control action need to be varied.

A possible solution to overcome this problem is to move from an embedded (or internal) control strategy, where the biomolecular networks are all embedded in a single cell, to a multicellular control strategy (see [10]) where the required functions are distributed across different cell populations in a microbial consortium [11]. This can indeed reduce metabolic loads and minimize unwanted effects, such as retroactivity, by physically separating the various components of the design among different cells [12], [13].

In this letter, we first present a multicellular realization of the PID controller inspired from the embedded single-cell solution in [7]. Then, we focus on the implementation of P and PI controllers within a microbial consortium comprising different cell populations communicating through appropriate orthogonal *quorum sensing* molecules. After presenting abstract biological implementations for each of the proposed strategies, we derive analytical conditions for tuning the controller gains, providing insights on the biological parameters that most influence their performance. We complement the theoretical derivations with a set of *in silico* experiments carried out using the realistic agent-based microbial simulator BSim [14], [15]. The results of all the experiments confirm the effectiveness of the proposed multicellular architectures whose *in vivo* implementation is the subject of ongoing research.

## II. Multicellular PID control strategy

We propose to realize a distributed biological PID controller entrusting each action to a different cellular population within a microbial consortium, see Fig. 1. Here, three cellular populations, denoted as *controllers*, implement the proportional, integral and derivative actions, respectively, implementing a biological equivalent of the classical PID control strategy given by:

**Fig. 1:**
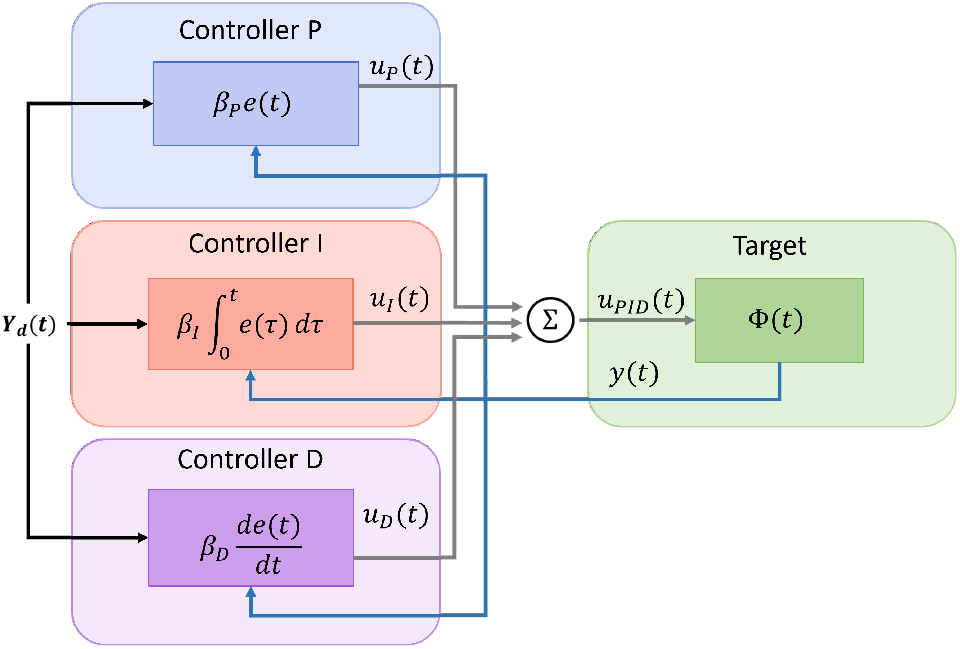
Schematic representation of a distributed biological PID controller. The three controllers compare the reference signal *Y*_d_(*t*) with the output of the target population *y*(*t*), and collectively compute the overall control signal *u*_*PID*_(*t*), closing the control loop and regulating the process Φ(*t*). Each controller realizes the biological equivalent of the classical PID control actions in (1).

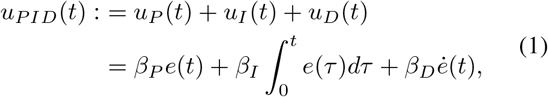

where *β*_*P*_, *β*_*I*_ and *β*_*D*_ play the role of the proportional, integral and derivative gains, and *e*(*t*) is the control error, which is a function of the measured output *y*(*t*) of the process Φ(*t*) under control and the desired value *Y*_d_(*t*). The overall control signal *u*_*PID*_(*t*) computed by the controllers is sensed by the *target* population hosting the process Φ(*t*), whose output *y*(*t*) is fed back to the controllers closing the control loop.

Here, we present two control architectures stemming from the full schematic in Fig. 1; the former composed by one population implementing the Proportional (P) action that controls the targets, and the latter in which an additional population is inserted in the consortium to implement an Integral (I) action (Fig. 2). For the sake of brevity, we leave the study of other possible configurations for a future work. The communication between controllers and targets is realized by the pair of orthogonal *quorum sensing* molecules *Q*_*u*_ and *Q*_*x*_ that act as proxies of the control input *u*_*PID*_(*t*) and the measurement of the process state *y*(*t*), respectively, that are produced by the cells and diffuse through their membranes into the environment.

**Fig. 2:**
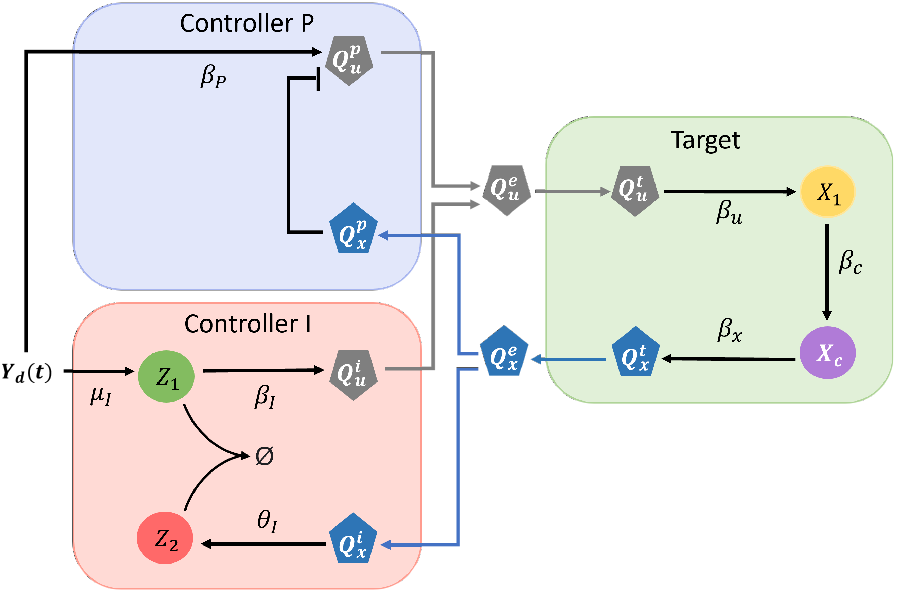
Abstract implementation of a distributed biological PI controller. The output of the process Φ(*t*) is the quorum sensing molecule *Q*_*x*_, produced proportionally to the target gene *X*_*c*_. Its input is the gene *X*_1_, which is actuated by the *control* quorum sensing molecule *Q*_*u*_. Each controller population evaluates the control error *e*(*t*) by comparing the reference signal *Y*_d_(*t*) and the process output carried by *Q*_*x*_, thus contributing to the overall production of *Q*_*u*_. Circles represent internal molecular species, while polygons represent the signaling molecules.

Each controller population senses the control error *e*(*t*) by comparing the reference signal *Y*_d_(*t*) with the measure of the output carried by the *sensing* quorum sensing molecule *Q*_*x*_. It then contributes accordingly to the overall production of the quorum sensing molecule *Q*_*u*_ which delivers the control input affecting the target cells. Therein, the process Φ(*t*) can be any network of genes that is directly affected by *Q*_*u*_ and whose output is the expression of some gene of interest. To close the loop, the process Φ(*t*) activates the production of *Q*_*x*_ that, by diffusing into the environment, can be sensed by the controllers.

Next, we derive the mathematical models of both schemes, describing the *aggregate* dynamics of the populations, that is, the evolution of the concentrations of the chemical species involved in the loop averaged over the entire consortium. We assume that all populations in the consortium are equally balanced, which also implies that the molecules diffuse through the cell membrane with the same diffusion rate *η*. This assumption is later relaxed in Section IV where *in silico* experiments are carried out to evaluate the impact of cell-to-cell variability and spatio-temporal effects on the control performance. Using the derived models, we analyze steady-state properties of each control strategy, deriving sufficient conditions on the control parameters that guarantee regulation and tunability of the biological process of interest. The superscripts *e, t, p, i*, are used in the rest of this letter to refer to quantities in the environment, in the target cells, in the Proportional, or in the Integral controller cells, respectively.

### A. Mathematical modelling

We assume that the output of the process Φ(*t*) is the quorum sensing molecule *Q*_*x*_, produced proportionally to the target gene *X*_*c*_, and that its input is the gene *X*_1_, which is actuated by *Q*_*u*_ (Fig. 2).

The dynamics of the network hosted in the target cells can be described by the following set of ODEs:

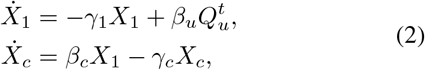

where *γ*_1_ and *γ*_*c*_ are the degradation rates of the species *X*_1_ and *X*_*c*_, respectively, and *β*_*c*_ and *β*_*u*_ are activation rates, modeling the strength of the activation induced by transcription factors *X*_1_ and *Q*_*u*_ [5], [7], [16]. The sensing molecule and process output *Q*_*x*_ is produced by the target cells as a function of the species *X*_*c*_ (for the sake of simplicity we assume the production of *Q*_*x*_ is proportional to *X*_*c*_). Hence, information about the target state is broadcast to the other cells in the consortium by *Q*_*x*_ diffusing into the environment with a dynamics assumed here to be linear. Under these assumptions, the dynamics of the sensing molecule in the targets can be modeled as:

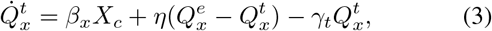

where *β*_*x*_ is the activation rate due to *X*_*c*_, *η* is the diffusion rate of the molecule *Q*_*x*_ across the cell membrane, and *γ*_*t*_ is the dilution rate of *Q*_*x*_ in the target cells.

The Proportional controller is implemented here as in [7] by means of an inhibitory action whose strength is proportional to the output fed back from the targets combined with an activation proportional to the amplitude of the reference signal. Namely, we can describe the dynamics of the intracellular concentration 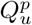 as:

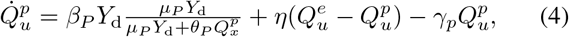

where *γ*_*p*_ is the dilution rate of *Q*_*u*_ inside the Proportional controller cells, *β*_*P*_ represents the maximal production rate of the control molecule and plays the role of the proportional gain as in (1), *μ*_*P*_ and *θ*_*P*_ are positive coefficients characterizing the control action [7]. It can be shown (see [7] for details) that the first term in (4) corresponds to an action that is proportional to the control error 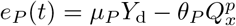.

To implement the Integral action, an antithetic motif [5] is embedded into the controller population. This module uses a pair of chemical species, say *Z*_1_ and *Z*_2_, produced proportionally to the reference signal *Y*_d_ and to the sensing molecule *Q*_*x*_, respectively, able to annihilate each other with a high affinity. The dynamics of the network embedded in this population can then be described by the set of ODEs:

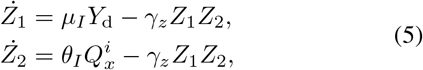

in which *μ*_*I*_*Y*_d_ and 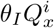 are production rates, *γ*_*z*_ is the annihilation rate between *Z*_1_ and *Z*_2_, which is assumed here to be the only source of degradation for the species *Z*_1_ and *Z*_2_, as also done in [7], [16]. Following similar arguments to those in [7], it can be shown that 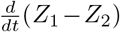 is proportional to the control error, defined as 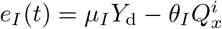.

The network described in (5) is complemented with the control molecule dynamics, described as:

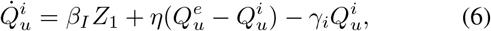

where *β*_*I*_ plays the role of the integral gain and all the other parameters have an analogous meaning as those in (3).

Finally, the mathematical models are completed by adding ODEs describing the diffusion dynamics of the quorum sensing molecules across the cell membranes. Specifically, the concentration of the molecules into the environment is described by:

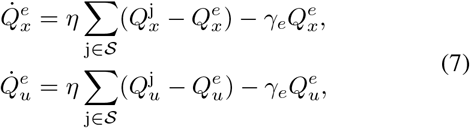

where 𝒮 = {*p, t*} if the Proportional controller population is only present, while 𝒮 = {*p, i, t*} when the Integral controller population is also added to the consortium. The dynamics of the concentrations of *Q*_*x*_ and *Q*_*u*_ inside those cells not directly producing them is given by:

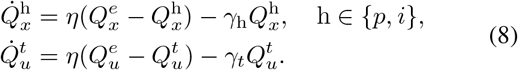

## III. Circuit Design

We derive some analytical conditions on the parameters of the genetic circuits which guarantee successful regulation of the measured output *Q*_*x*_ to the desired value. To this aim, we first derive a reduced order model of the consortium dynamics and then, via a stability analysis, we provide sufficient conditions that the biomolecular parameters must satisfy in order for the consortium to operate correctly.

### A. Assumptions and problem statement

To derive simple, yet meaningful, analytical conditions guiding the design of the controller populations, we make some realistic assumptions on the values of the parameters. Namely, we assume that (i) each population divides at the same rate, implying that the dilution rates for all species are identical (i.e. *γ*_*p*_ = *γ*_*i*_ = *γ*_*t*_ = *γ*_1_ = *γ*_*c*_ = *γ*); (ii) the degradation of each quorum sensing molecule in the external growth environment can be neglected, i.e. *γ*_*e*_ = 0; and (iii) the promoters induced by the reference signal *Y*_d_ and the sensing molecule *Q*_*x*_ are the same in the Proportional and Integral controllers, implying that *μ*_*p*_ = *μ*_*i*_ = *μ* and *θ*_*p*_ = *θ*_*i*_ = *θ*. Also, we further assume that:

#### Assumption 1

*The quorum sensing molecules diffuse faster than they degrade*.

#### Assumption 2

*The annihilation process between the species Z*_1_ *and Z*_2_ *is fast enough*.

Defining Γ_*P*_ := *γ*_*t*_ + *γ*_*p*_ + *γ*_*e*_ = 2*γ* and Γ_*PI*_ := *γ*_*t*_ + *γ*_*p*_ + *γ*_*i*_ +*γ*_*e*_ = 3*γ*, Assumption 1 translates to *η*≫ Γ_*P*_ when only Proportional controller cells are present, and to *η* ≫Γ_*PI*_ when both controller populations are present. Assumption 2 implies that 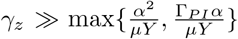, where *α* is a function of the model parameters defined as

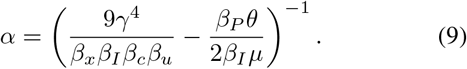

Given the assumptions above and defining the control error as:

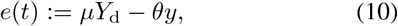

where we defined the controlled output as 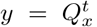, the control problem is that of engineering a microbial consortium comprising one or more controller populations so that at steady state the output of the process in the target cells is robustly regulated to the desired value, that is, lim_*t*→∞_ *e*(*t*) = 0.

### B. Multicellular P controller

Under the assumptions made in Section III-A and quasi-steady state assumption on the quorum sensing dynamics, when the targets are regulated by controllers solely implementing a Proportional control action, the dynamics of the consortium can be approximated by the following reduced order model:

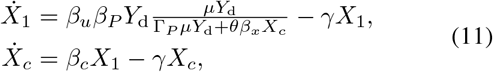

where the controlled output is defined as 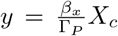 (see Appendix A for details). It can be demonstrated that model (11) has a unique admissible equilibrium point, which is always locally asymptotically stable. However, for the sake of brevity, the proof is omitted here. Moreover the steady-state error *e*_ss_ is given as:

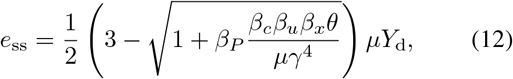

which nonlinearly depends on the value of the proportional gain *β*_*P*_ and that can be made closer to zero by selecting *β*_*P*_ as:

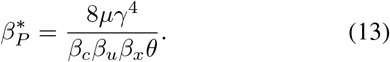

Indeed, if it were possible to select 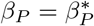, we could have 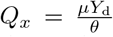 at steady state, which implies *e*(*t*) = 0. Note that such a choice would require perfect knowledge of the target cells parameters which is unrealistic. We will therefore explore later in Section IV how the error varies for values of *β*_*P*_ in a given range of interest and evaluate the effects of *β*_*P*_ on the dynamics of 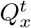, assessing the sensitivity of the control strategy to parameter mismatches or uncertainties.

### C. Multicellular PI controller

A possible solution to overcome model uncertainties and robustify the designed control system is to add a third population implementing an Integral control action. Under the same assumptions made in Section III-A, the dynamics of the resulting consortium comprising both the P and I controller populations can be approximated by the following set of ODEs:

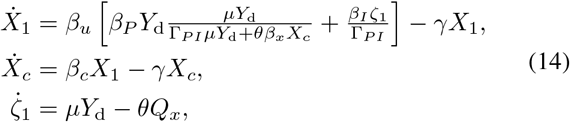

where *ζ*_1_ = *Z*_1_ − *Z*_2_ and 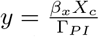. Details on the derivation of equation (14) are reported in Appendix A. This dynamical system has a unique, non-negative equilibrium point if the proportional gain is chosen such that:

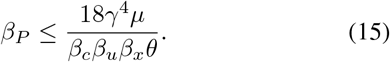

Reaching this equilibrium point ensures that 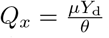, thus *e*(*t*) = 0. By carrying out a local stability analysis, we found that the equilibrium point is locally asymptotically stable if the value of the integral gain *β*_*I*_ does not exceed a threshold dependent on the other parameters including the proportional gain *β*_*P*_, that is:

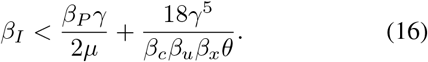

Condition (16) gives insights on the biological elements that influence the performance of the PI multicellular architecture. Specifically, choosing fast dividing cells (i.e. high values for *γ*) or reducing of the strength of the promoters induced by the reference *Y*_d_ and the sensor molecule *Q*_*x*_ can widen the range of values of *β*_*I*_ that guarantee the correct operation of the control consortium.

## IV. In silico control experiments

To validate the proposed multicellular control architectures we carried out *in silico* experiments in BSim [14], [15], an agent-based environment explicitly designed to simulate bacterial populations. BSim allows to keep track of each cell in the consortium, simulating both the dynamical processes hosted in the organism and the bio-mechanics of the cell. In addition, the diffusion of the quorum sensing molecules, cell growth and division, and cell-to-cell variability can be explicitly simulated together with realistic geometric constraints of the host environment. In particular, BSim accurately simulates cells cultured in a micro-environment such as a microfluidic device where nutrients are constantly provided, allowing cells to grow in exponential phase. Here, we emulated a scaled version of the device described in [17], [18], that is, a microfluidic chamber of dimensions 17 *μ*m × 15 *μ*m × 1 *μ*m, which can host around 100 cells. These dimensions were selected as a good trade-off between number of cells hosted and computational burden. The growth and mechanical parameters were selected as in [11], while the nominal values of the parameters in the network were chosen as described in Appendix B.

Selecting 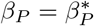, both control architectures showed good regulation capabilities of the output species 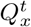 over a period of 12000 min at different set-point values (Fig. 3, blue curves), with a settling time of about 500 min for the Proportional controller (Fig. 3a) and about 700 min for the PI controller (Fig. 3b). In all simulations *β*_*I*_ was selected according to (16). However, when *β*_*P*_ cannot be tuned to match equation (13), the Proportional controller alone fails to regulate *y* to the desired value, showing increasingly higher value of the steady-state errors as *β*_*P*_ decreases. Instead, as expected, adding the Integral control action, the steady-state error is not sensitive to changes in the value of *β*_*P*_, as long as (15)-(16) are satisfied.

**Fig. 3:**
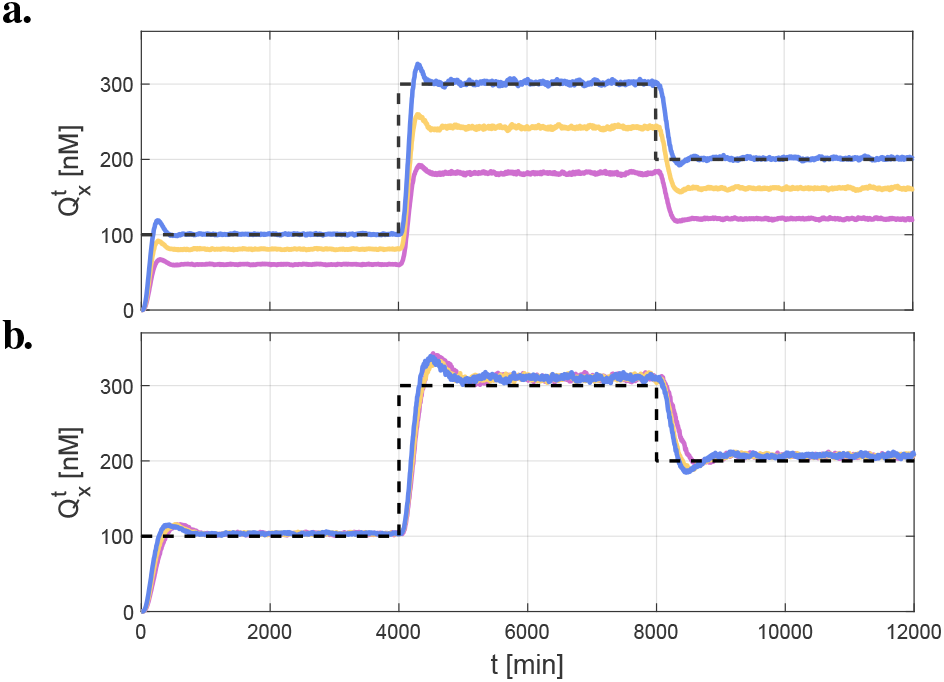
Set-point tracking experiments in BSim: evolution in time of the average concentration in the targets of the quorum sensing molecule 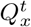 when they are controlled by the Proportional controllers only (panel a) and by a PI control action (panel b). The control gains were selected as *β*_*P*_ ={0.02, 0.03, 0.04} min^−1^ (purple, yellow and blue line, respectively) and *β*_*I*_ = 0.0002 min^−1^. The piece-wise constant reference signal *μY*_d_(*t*)*/θ* is depicted as a dashed line.

Next, we tested the robustness of the control architectures with respect to imbalances in the relative composition of the consortium. Indeed, despite the host cells being identical, unavoidable asymmetries in the metabolic load on each population can cause their relative numbers to change over time. To this aim, we varied the populations’ relative numbers defined as 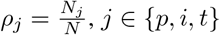, where *N*_*j*_ is the number of individuals belonging to the *j*-th population, while keeping constant the total number *N* of cells in the chamber, and evaluating the percentage error at steady state defined as:

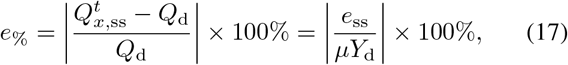

where *e*_ss_ is the control error (10) at steady state, 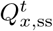 is the value of 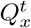 at steady state, and 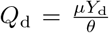. Note that keeping *N* constant implies that *ρ*_*p*_ + *ρ*_*i*_ + *ρ*_*t*_ = 1.

Fig. 4 shows that the Proportional controller works best when the populations are close to balance, while exhibiting increasingly higher errors at steady state as imbalance between controllers and targets increases, with a maximum error of 70% when the imbalance is extreme (*ρ*_*p*_ *<* 0.1). Adding an Integral contribution to the control action significantly increases the architecture performance and robustness, with the relative error never exceeding 30% even when the imbalance of the controllers and target populations’ numbers are consistent.

**Fig. 4:**
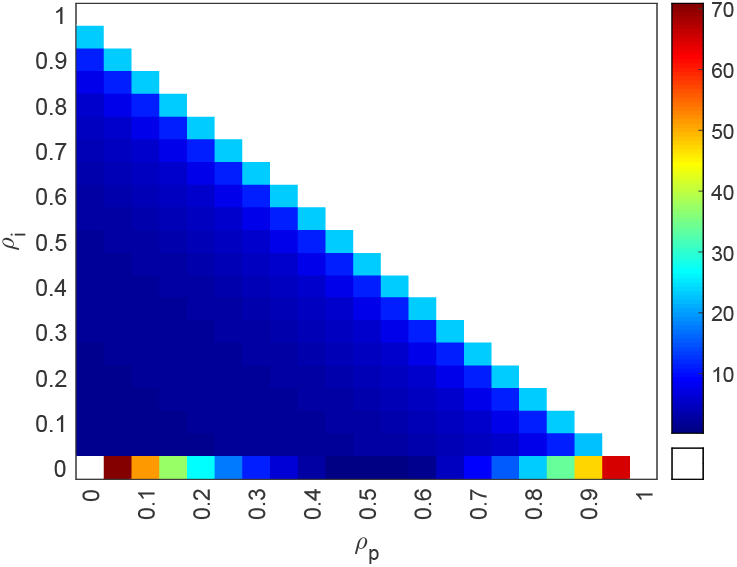
Robustness to imbalances in the consortium composition: percentage error at steady state (17) as the relative ratios of the three populations are changed. The total number of cells is fixed in BSim to *N* = 20, with no growth dynamics and the ratio *ρ*_*j*_ for *j* ∈ {*p, i, t*} is constrained to *ρ*_*p*_ + *ρ*_*i*_ + *ρ*_*t*_ = 1. The reference signal is fixed to *Y*_d_ = 60 nM, while the gains are chosen as *β*_*P*_ = 0.0414 min^−1^ and *β*_*I*_ = 0.0002 min^−1^.

Finally, we tested the effects of cell-to-cell variability on the overall control performance. This heterogeneity in the response of the cells is mainly due to variations between plasmid copy numbers among different individuals in a population, caused by a possibly uneven distribution of the genetic material between cells after division. To model this effect, at each cell division, the value of all parameters of the daughter cells were drawn from a normal distribution centered at their nominal values *μ* with standard deviation *σ* = *CV · μ*, where *CV* is the coefficient of variation. The sensitivity of the control system was evaluated, as *CV* increases, by computing the percentage error at steady state, defined as:

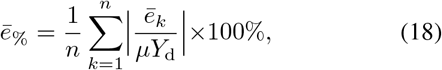

where *ē*_*k*_ is the control error (10) averaged over the last 5000 minutes in the *k*-th experiment, and *n* is the total number of experiments. We observed that both the P and the PI control architectures guarantee small sensitivity to increasing levels of heterogeneity within the cellular populations with the relative error never exceeding 10% (Fig. 5). However, the multicellular PI control architecture shows higher robustness with the steady-state error showing much smaller variations under perturbation.

**Fig. 5:**
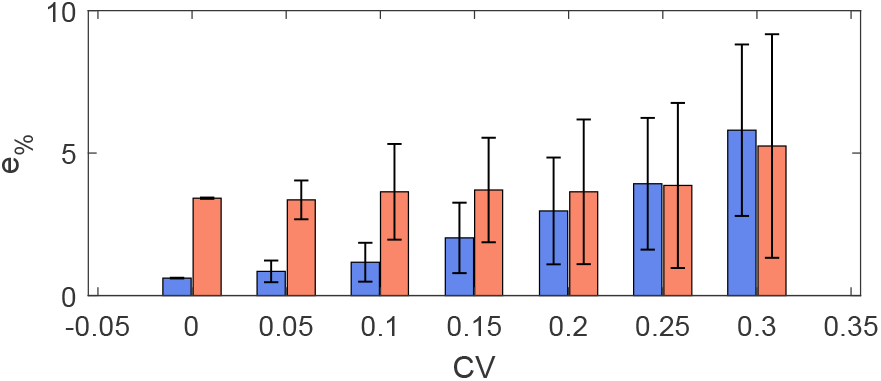
Robustness to cell-to-cell variability: average percentage error at steady state and standard deviation as the heterogeneity of the parameters increases, when the consortium is regulated by P controllers only (blue bars) and PI controllers (orange bars). For each value of *CV* ∈ {0, 0.05, 0.1, 0.15, 0.2, 0.25, 0.3} we performed *n* = 50 simulations drawing independently all cells’ parameters from a normal distribution centered at their nominal value *μ* with standard deviation *σ* = *CV · μ*. The reference signal is fixed to *Y*_d_ = 60 nM, while the gains are chosen as *β*_*P*_ = 0.0414 min^−1^ and *β*_*I*_ = 0.0002 min^−1^. All simulations were performed for a total time of 8000 min, using a chamber of dimensions 5.7 *μ*m × 15 *μ*m × 1 *μ*m.

## V. Conclusions

We investigated first analytically and then numerically two multicellular architectures where a process of interest hosted in a target cell population was regulated using a P or a PI control law implemented across other populations in the consortium. We showed that it is possible to choose the control gains so that successful regulation of the controlled protein in the target cells at the desired level is achieved. In addition, we observed that, similarly as in embedded antithetic controllers, perfect robust adaptation can only be achieved when an additional population implementing an integral control action is included in the consortium. Finally, we validated the effectiveness and robustness of the designed consortia via realistic agent-based simulations in BSim showing their viability even in the presence of unavoidable effects due to cell-to-cell variability, diffusion and cell growth.

A key open problem towards the *in vivo* implementation of the proposed multicellular PI architecture is to guarantee stable co-existence and maintain a desired ratio between the cell populations involved. This is possible either by embedding in the cells extra gene pathways to regulate the growth rates of the populations, or by culturing the cells in a controlled environment where it is possible, by means of an external control action, to guarantee that the relative numbers of the populations within the consortium are kept within acceptable bounds [18]–[21].

## Appendix

### A. Derivation of the reduced order models

Under the conditions presented in Section III-A, the dynamics of the microbial consortium comprising three populations and described by equations (2)-(8) can be further simplified making a quasi-steady state assumption on the dynamics of the quorum sensing molecules, that is, by imposing 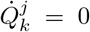, with *k* ∈ *{u, x}* and *j* ∈ *{t, p, i, e}*. Under Assumption 1, substituting their steady-state values in 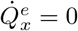 and 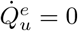 we obtain that:

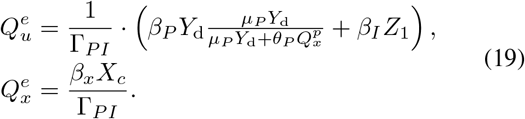

Then, by substituting (19) in 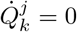, where *k* ∈ *{x, u}* and *j* ∈ *{p, i, t}*, and leveraging again that *η* ≫ Γ_*PI*_, we obtain 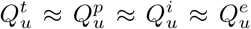, and 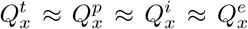. We then define those two quantities as *Q*_*u*_ and *Q*_*x*_ and substitute their steady state values in equations (2) and (5).

In addition, we introduce the change of variables *ζ*_1_ = *Z*_1_ − *Z*_2_, *ζ*_2_ = *Z*_2_, transforming equations (5) in:

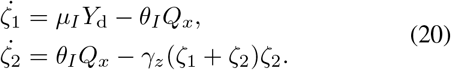

As done in [22], we can then use time scale separation on equations (20) to reduce the model to 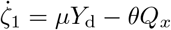.

The same procedure can be repeated, excluding the time scale separation operated on *ζ*_1_, *ζ*_2_, to retrieve a reduced order model for a consortium without the Integral controller population. The results are the same, with the exception of equation (19), in which Γ_*P*_ is in place of Γ_*PI*_, and the term *β*_*I*_*Z*_1_ is not present.

### B. Nominal biochemical parameters

The nominal biochemical parameters used in the BSim simulations are chosen as: *β*_*u*_ = 0.06 min^−1^, *β*_*x*_ = 0.03 min^−1^, *γ* = 0.023 min^−1^, *η* = 2 min^−1^ (taken from [11]); *β*_*c*_ = 0.1 min^−1^, *μ* = 1 min^−1^, *θ* = 0.3 min^−1^, *γ*_*z*_ = 0.01 nM^−1^min^−1^ (taken from [7]).

